# The β3-integrin endothelial adhesome regulates microtubule dependent cell migration

**DOI:** 10.1101/145839

**Authors:** Samuel J Atkinson, Aleksander M Gontarczyk, Tim S Ellison, Robert T Johnson, Benjamin M Kirkup, Abdullah Alghamdi, Wesley J Fowler, Bernardo C Silva, Jochen J Schneider, Katherine N Weilbaecher, Mette M Mogensen, Mark D Bass, Dylan R Edwards, Stephen D Robinson

**Affiliations:** School of Biological Sciences, University of East Anglia, Norwich, UK.; Luxembourg Center for Systems Biomedicine (LCSB), University of Luxembourg, Luxembourg & Saarland University Medical Center, Internal Medicine II, Homburg, Germany.; Department of Internal Medicine, Division of Molecular Oncology, Washington University in St Louis, St. Louis, MO, USA.; Centre for Membrane Interactions and Dynamics, Department of Biomedical Science, University of Sheffield, Sheffield, UK.; Faculty of Medicine and Health, University of East Anglia, Norwich, UK.

## Abstract

Integrin β3 is seen as a key anti-angiogenic target for cancer treatment due to its expression on neovasculature, but the role it plays in the process is complex; whether it is pro- or anti-angiogenic depends on the context in which it is expressed. To understand precisely β3’s role in regulating integrin adhesion complexes in endothelial cells, we characterised, by mass spectrometry, the β3- dependent adhesome. We show that depletion of β3-integrin in this cell type leads to changes in microtubule behaviour that control their migration. β3- integrin regulates microtubule stability in endothelial cells through Rcc2/Anxa2 driven control of Rac1 activity. Our findings reveal that angiogenic processes, both *in vitro* and *in vivo*, are more sensitive to microtubule targeting agents when β3-integrin levels are reduced.

## INTRODUCTION

Angiogenesis, the formation of new blood vessels from those that already exist, plays an essential role in tumour growth (Hanahan and Weinberg, 2011). As such, targeting angiogenesis is seen as crucial in many anti-cancer strategies (Zhao and Adjei, 2015). Therapies directed against vascular endothelial growth factor (VEGF) and its major receptor, VEGF-receptor-2 (VEGFR2), whilst effective in a number of cancers, are not without side-effects due to the role this signaling pathway plays in vascular homeostasis (Chen and Hung, 2013). Fibronectin (FN)-binding endothelial integrins, especially αvβ3- and α5β1-integrins, have emerged as alternative anti-angiogenic targets because of their expression in neovasculature (Brooks et al., 1994; Kim et al., 2000). However, neither global nor conditional knockouts of these integrins block tumour angiogenesis long-term (Murphy et al., 2015; Reynolds et al., 2002; Steri et al., 2014), and clinical trials of blocking antibodies and peptides directed against these extracellular matrix (ECM) receptors have been disappointing (Schaffner et al., 2013; Stupp et al., 2014).

To gain novel insight into how αvβ3-integrin regulates outside-in signal transmission (Hynes, 2002), we have undertaken an unbiased analysis of the molecular composition of the mature endothelial adhesome (the network of structural and signaling proteins involved in regulating cell-matrix adhesions (Zaidel-Bar et al., 2007)), and profiled changes that occur when β3-integrin function or expression are manipulated**. In so** doing, we have uncovered β3-integrin dependent changes in microtubule behaviour that regulate cell migration.

## RESULTS AND DISCUSSION

The isolation and analysis of integrin adhesion complexes (IACs) by mass-spectrometry (MS) is difficult because of the low affinity and transient nature of the molecular interactions occurring at these sites. However, using cell-permeant chemical crosslinkers improves recovery of IAC proteins bound to either FN-coated microbeads (Horton et al., 2015) or plastic dishes(Schiller et al., 2011). These advances have led to the characterisation of IACs from a number of cell types. Whilst a core consensus adhesome (the network of structural and signaling proteins involved in regulating cell-matrix adhesion (Zaidel-Bar et al., 2007)) can be defined (Horton et al., 2015), the composition and stoichiometry of the meta-adhesome depends on the cell-type being analysed, the integrin-receptor repertoire expressed by that cell type, and on any imposed experimental conditions.

To examine the composition of the endothelial adhesome we isolated lung microvascular endothelial cells (ECs) from C57BL6/129Sv mixed background mice and immortalised them with polyoma-middle-T-antigen by retroviral transduction (May et al., 2005). As our main interest was in establishing how β3-integrin influences the endothelial adhesome, we adhered cells to FN for 90 minutes, which allows β3-rich (mature) focal adhesions (FAs) to form (Schiller et al., 2013). To distinguish integrin-mediated recruitment of proteins from non-specific background, we also plated cells on poly-L-lysine (PLL) as a negative control (adhesion to PLL does not depend on integrins). Visualisation of neuropilin-1 staining in whole cells showed that this protein, which we previously demonstrated is present in the mature EC adhesome (Ellison et al., 2015), co-localises with talin-1 in FAs when cells are plated on FN, but not PLL (Fig. 1a). For all proteomics experiments, we crosslinked FAs using the cell permeant and reversible cross-linkers DPDPB and DSP (see materials and methods) for 5 minutes. Cells were lysed and subjected to a high sheer flow water wash to remove non-crosslinked material. Crosslinking was reversed, and samples were precipitated and eluted for analyses. Prior to MS, samples were quality controlled by SDS/PAGE and silver-staining to ensure efficient removal of non-crosslinked material had occurred (Fig. 1b).

**Figure 1.**
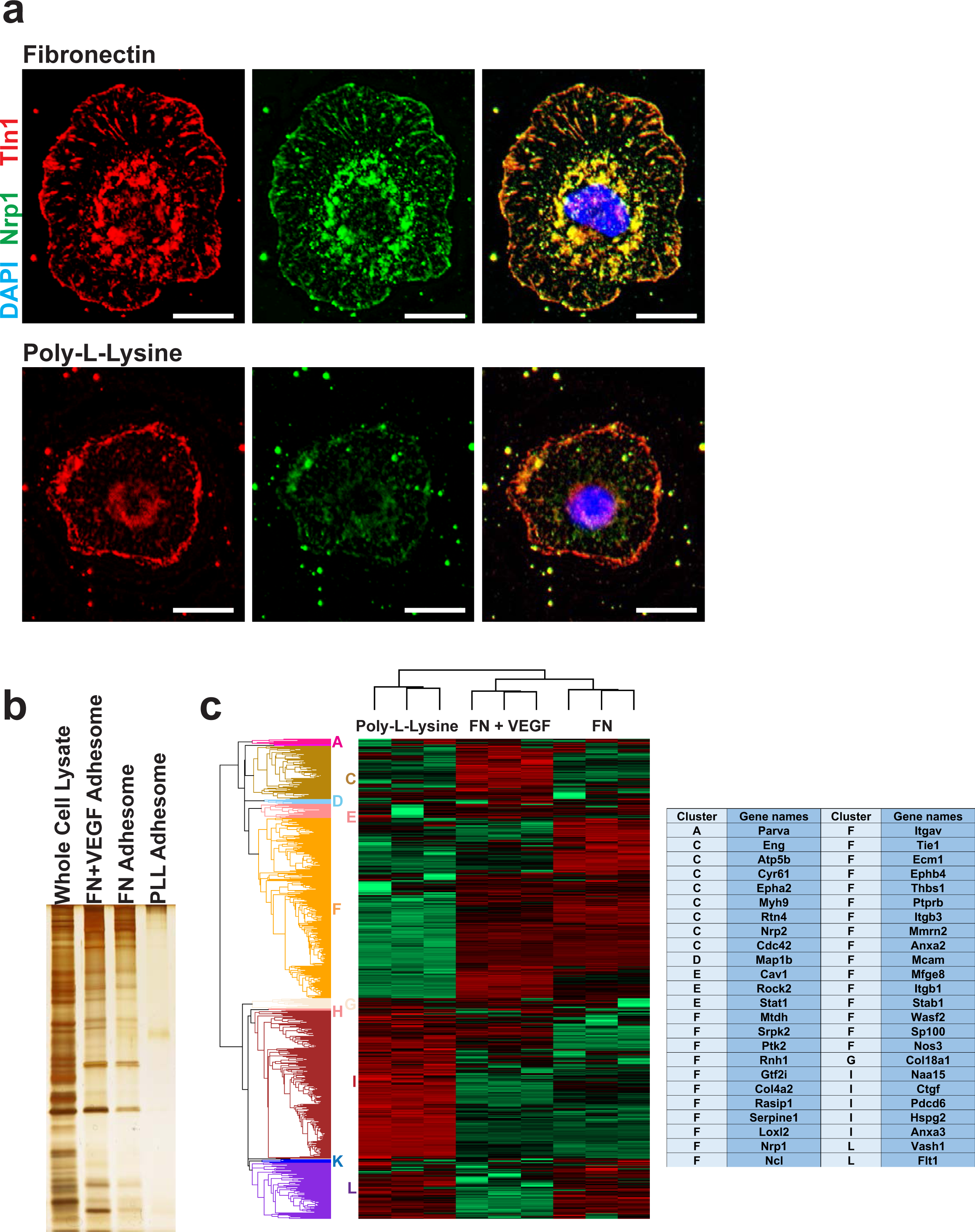
Defining the FN endothelial adhesome. **a)** WT ECs were adhered to fibronectin (top row) or poly-l-lysine (bottom row) coated coverslips for 90 minutes before fixing and immunostaining for neuropilin-1 (Nrp1-green) and talin-1 (Tln1-red) along with a nuclear stain (DAPI-blue). Scale bar = 10 μm. **b)** An example silver stain used in the quality control of adhesome samples. Adhesome enrichment was carried out on 6 × 10^6^ WT ECs on fibronectin (FN), fibronectin with VEGF (FN + VEGF) or poly-l-lysine (PLL) before acetone precipitation. After resuspension samples were run on a SDS-PAGE gel along with a whole cell lysate control and silver stained. **c)** Triplicate adhesome samples from WT ECs adhered on fibronectin (FN), fibronectin with VEGF (FN + VEGF) or poly-l-lysine were sent for quantitative mass spec analysis. Label free quantification was carried out using MaxQuant followed by analysis in Perseus. Unsupervised hierarchical clustering (Euclidian distance calculation) was carried out with red showing highly abundant proteins and green showing low abundance proteins. 12 significant clusters were automatically identified using a distance threshold of 3.34 and labelled A-L. Angiogenesis associated proteins were identified using GOBP annotations (GO:0001525, GO:0002040, GO:0002042, GO:0016525, GO:0045765, GO:0045766) and are displayed in the table along with their associated cluster.

Label-free proteomic analyses of the FN + VEGF, FN, and PLL adhesomes (Fig. 1c; Supplementary Table SI) initially detected and quantified 1468 proteins. Stringent filtering, requiring proteins to be detected in all 3 repeats of at least one condition, left 1064 proteins - a high confidence dataset that was used to define the endothelial adhesome. Hierarchical clustering based on average Euclidian distance identified 12 clusters (A-L) which could be considered VEGF-enriched proteins (A-C), FN-enriched (D-F), and PLL-enriched (G-L). Fisher’s exact test enrichment analysis was carried out to identify which pathway, process, or component proteins within these clusters belong to using Gene Ontology annotations. Cell projection (GOCC, p=8.62 × 10^−5^) and microtubule (GOCC, p=1.6 × 10^−4^) categories were significantly enriched when cells were treated with growth factor, suggesting they are important in VEGF-mediated processes. Leukocyte trans-endothelial migration (KEGG, p=9.71 × 10^−5^) proteins were enriched in the FN adhesome, but not in the VEGF-stimulated adhesome, suggesting our cells represent quiescent vasculature without VEGF-stimulation. This same category contains many endothelial specific proteins (e.g. ve-cadherin/Cdh5), further confirming that the cells have an endothelial identity. Focal adhesion (KEGG, p=9.31x10^−7^) proteins were enriched in the FN adhesome but depleted in the PLL adhesome, confirming the success of the adhesome enrichment process, MS, and downstream analysis. Other adhesion/migration associated categories: focal adhesion (GOCC, p=5.99 × 10^−5^), cell projection (GOCC, p=3.03 × 10^−5^), cell adhesion (GOBP, p=1.61 × 10^−6^) and lamellipodium (GOCC, p=1.38 × 10^−4^) were depleted in the PLL adhesome.

FN-based matrices are essential for angiogenesis (Zhou et al., 2008) and studies in fibroblasts have demonstrated that αv-integrins and α5β1-integrin cooperate to direct cell migration on FN-based matrices(Schiller et al., 2013). Therefore, we wanted to profile changes in the FN-endothelial adhesome in response to αvβ3-integrin blockade. Thus, we first determined the adhesome composition of ECs treated with EMD66203 (cyclo-Arg-Gly-Asp-DPhe-Val), a conformationally constrained cyclic pentapeptide that selectively targets αvβ3-integrin (Pfaff et al., 1994). EMD66203 inhibited EC adhesion to FN (the effects of EMD66203 on adhesion to vitronectin, an αvβ3 specific ligand, was tested to ensure activity of the compound - Fig. 2a) but had no dramatic effect on the endothelial adhesome (Fig. 2b; Supplementary Table S2), suggesting that: (1) even if αvβ3-integrin is unable to bind ECM, it can localise to FAs and participate in adhesome assembly in the presence of EMD66203; or (2) under the experimental conditions used, only a small percentage of αvβ3-integrin is conformationally available for EMD66203 binding (Demircioglu and Hodivala-Dilke, 2016).

**Figure 2.**
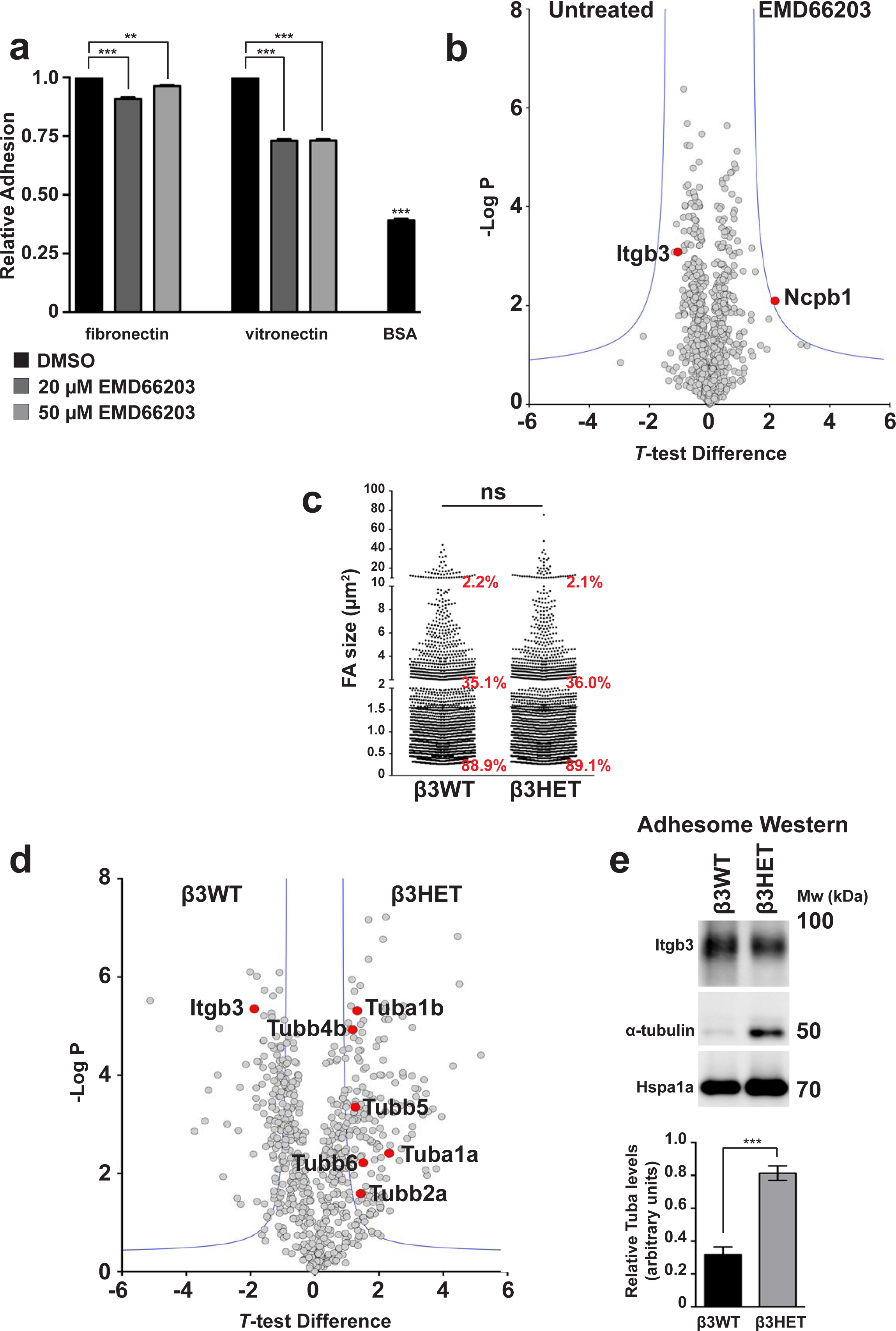
Analysis of the β3-integrin dependent adhesome. **a)** Adhesion analysis of endothelial cells adhered to saturating concentrations of fibronectin or vitronectin in the presence of EMD66203, an αvβ3-integrin specific RGD mimetic. Bars = mean (±SEM) adhesion relative to vehicle control (DMSO). **b)** Visual representation of the significant analysis of microarrays (SAM) method as a volcano plot for DMSO versus 20μM EMD66203 treated endothelial cell samples (n=3). T-test difference is plotted against 2013;log of the P value. The blue lines show the cut-off for significance as defined by the SAM. Integrin β3 (Itgb3) as well as Ncpb1 (the only significant change) have been highlighted as red points. **c)** Distribution of adhesion size classes (0-2μm, 2-10μm; >10μm) in β3WT versus β3HET endothelial cells (n>1400 FAs per genotype). **d)** Visual representation of the significant analysis of microarrays (SAM) method as a volcano plot for β3WT and β3HET samples (n=3). T-test difference is plotted against –log of the P value. The blue lines show the cut-off for significance as defined by the SAM. Integrin-β3 (Itgb3) as well as all detected tubulins (Tub) have been highlighted as red points. **e)** Adhesome samples from β3WT and β3HET endothelial cells adhered to fibronectin. Samples were Western blotted for integrin-β3 (Itgb3), α-tubulin and heat shock protein 70 (Hspa1a). Blot shown is representative of the 5 individual experiments that are quantified in the bar graph below. Bars = mean (±SEM) relative α-tubulin levels normalised to Hspa1a levels. ns = not significant; **= P<0.01; ***= P<0.001 in an unpaired, two-tailed t-test.

To test the consequences of excluding β3-integrin more efficiently from the EC adhesome, we decided to profile changes in β3-heterozygous (β3HET) ECs, which carry one wild-type allele of β3-integrin, and one knockout allele. These cells express 50% wild-type levels of β3-integrin and we have shown they are a good model for studying the role of αvβ3-integrin in cell migration, whilst evading changes arising from the complete loss of the integrin on both alleles (e.g. altered VEGFR2-mediated responses) (Ellison et al., 2015). Both wild-type (β3WT) and β3HET ECs adhere equally to saturating concentrations (10 μg/ml) of FN (see Ellison *et al.*, 2015 (Ellison et al., 2015)). To compare the size distribution of FAs between β3WT and β3HET ECs (which might affect the stoichiometry of components in the adhesome), we seeded cells for 90 minutes on FN, immunostained for paxillin, and measured FA area; we noted no differences in the percentage of FA size distributions between the two genotypes (Fig. 2c). Therefore, MS analyses comparing the adhesome between β3WT and β3HET ECs were performed (Fig. 2d; Supplementary Table S3). Enrichment analysis showed a depletion of cytoskeletal components (GOCC, p = 4.73 × 10^−5^) in the β3WT adhesome when compared with the β3HET adhesome, despite the enrichment of adhesion/migration associated categories previously noted in the FN adhesome of β3WT ECs (Fig. 1c). Whilst a majority of individual FA components in the mature adhesome do not change upon β3-integrin depletion, downstream connections to cytoskeletal components do. We took a particular interest in microtubules (MTs) because all detected tubulins were significantly upregulated in the β3HET adhesome. To confirm this finding by other means, we probed Western blots for α-tubulin and showed a significant increase in FA-enriched samples from β3HET cells compared with β3WT cells (Fig. 2e).

Our findings intimated that αvβ3-integrin drives MT localisation away from FAs. To increase the power of our studies, we included β3-integrin knockout (β3NULL) ECs in subsequent analyses. We examined MT organisation in β3WT, β3HET, and β3NULL ECs by immunolabeling for α-tubulin in whole cells (Fig. 3a). Although no gross changes in cell microtubule arrays were observed, the increased bundling of microtubules in the β3HET and β3NULL ECs suggests they could be crosslinked by microtubule stabilising proteins. Microtubule bundles are known to form in ECs to assist with directional migration by counteracting actomyosin generated contractile forces; these bundles often target adhesion sites in EC protrusions (Lyle et al., 2012). Furthermore, total cellular levels of α-tubulin were similar across the three genotypes (Fig. 3b). However, colocalisation of MTs with talin-1 at peripheral FAs was greater in β3HET and β3NULL ECs as was extension into lamellipodea (Fig. 3c). Overall, the findings suggest that β3-integrin limits the targeting of MTs to FAs.

**Figure 3.**
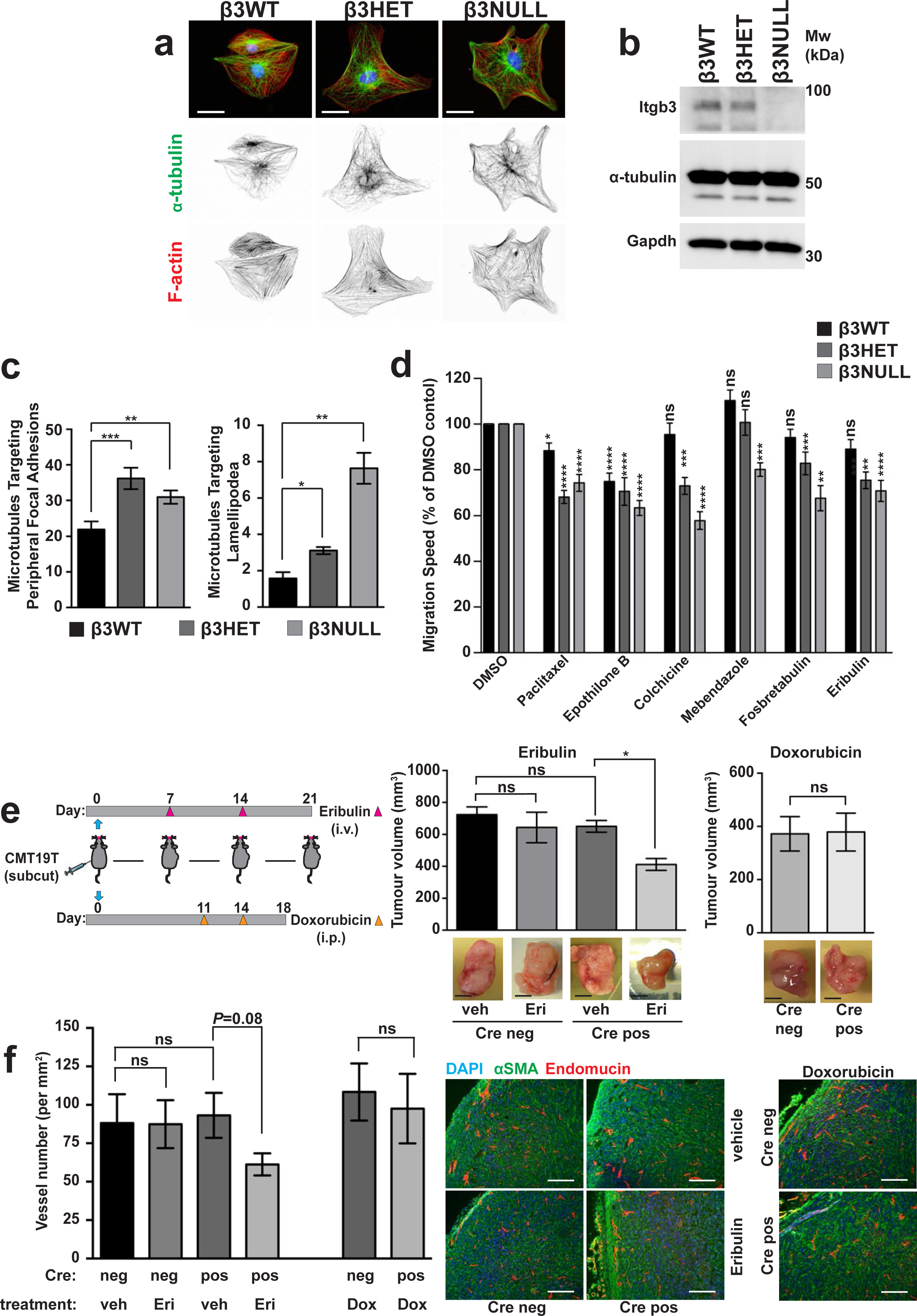
Analysis of microtubules in β3WT, β3HET and β3NULL endothelial cells. **a)** β3WT, β3HET and β3NULL endothelial cells were adhered to fibronectin coated coverslips for 90 minutes before being PHEMO fixed and immunostained for a-tubulin (green). Nuclear (DAPI-blue) and Phallodin (F-actin - red) stains were also used. Inverted black and white images of α-tubulin and F-actin are shown below the three-colour overlays. Scale bar = 10μm. **b)** β3WT, β3HET and β3NULL endothelial cells were adhered to fibronectin for 90 minutes before being lysed and Western blotted for Integrin-β3 (Itgb3), α-tubulin and Gapdh (as a loading control). **c)** *Left* β3WT, β3HET and β3NULL endothelial cells were adhered to fibronectin coated coverslips for 90 minutes before being methanol (-20°C) fixed and immunostained for α-tubulin and talin-1. The number of microtubules that terminated at a focal adhesion were counted for each genotype (n=15). *Right* β3 WT, HET and NULL ECS were transfected with paxillin-GFP and left to recover overnight. The cells were then adhered to fibronectin coated coverslips and allowed to recover for 3 hours before being treated with 100 nM SiRTubulin and 1 μM verapamil overnight. The next day, fresh media containing SiRTubulin and verampamil (same dose) was added cells were imaged every minute for 30 minutes (n=3). Areas of adhesive fronts were assessed by measuring the growth of paxillin-GFP positive areas between the 1st and 30th image. The number of microtubules that entered the adhesive front was quantified to give the number of microtubules entering lamellipodia relative to the area of adhesive fronts for each cell. **d)** β3WT, β3HET and β3NULL endothelial cells were adhered to fibronectin overnight. Migration speed of individual cells was measured over 15 hours using the MTrackJ plugin for ImageJ whilst under the influence of the indicated MTA. Migration speeds are shown as a percentage of the speed of the corresponding genotype under DMSO (vehicle) treatment (n>46). **e)** β3flox/flox Tie1Cre positive (pos) and negative (neg) animals were injected subcutaneously with 1X10^6^ CMT19T lung carcinoma cells and then treated as indicated in the schematic. Bar graph shows mean (± SEM) tumour volumes at the end of the experiment. Micrographs (below) show representative tumours from 2 independent experiments (n>5). Scale bars = 5mm. **f)** After excision, tumours from β3flox/flox Tie1Cre positive (pos) and negative (neg) animals were processed and endomucin staining was assessed over entire tumour sections to measure vascular density. Bars = mean (±SEM) vessel number per mm^2^ (n>5). Micrographs (right) show representative images of sections stained for alpha smooth muscle actin (αSMA=green), Endomucin (red) DAPI (blue). Scale bars = 100μm. ns = not significant; *= P<0.05; **= P<0.01; ***= P<0.001; ****= P<0.0001 in an unpaired, two-tailed t-test.

Given that MTs can drive FA turnover, and thus cell migration(Kaverina and Straube, 2011), we next tested whether EC migration is differentially sensitive to microtubule targeting agents (MTAs) in β3HET and β3NULL ECs. For each MTA examined, we first determined the dose of the compound that allowed 90 percent survival of β3WT ECs (not shown), and then tested the effects of this dose on random migration in β3WT, β3HET, and β3NULL cells (Fig. 3d). Random migration was affected by MT stabilisers (Paclitaxel, Epothilone B) in cells of all three genotypes, although β3WT cells were generally less sensitive than β3HET and β3NULL cells. β3WT ECs were insensitive to MT destabilisers (Colchicine, Mebendazole, Fosbretabulin) and the mechanistically unique MTA Eribulin (which functions through an end poisoning mechanism (Jordan et al., 2005)), whilst β3HET and β3NULL ECs showed a greater sensitivity to these compounds; in general, β3NULL cells were more sensitive than β3HET cells. We extended these types of analyses *in vivo* to examine the effects of Eribulin on tumour growth and angiogenesis. We chose Eribulin because the agent is well tolerated in mice (Su et al., 2016). We settled on a suboptimal dose (0.15 mg/kg) that would allow us to observe potential synergy with endothelial depletion of β3- integrin. β3-integrin-floxed/floxed mice (Morgan et al., 2010) were bred with Tie1Cre mice (Gustafsson et al., 2001) to generate β3-integrin-floxed/floxed Cre-positive animals (Cre-negative littermates were used as controls). CMT19T lung carcinoma cells were injected subcutaneously and allowed to establish for 7 days, at which point Eribulin was administered i.v. One week later, a second dose of Eribulin was given, and then tumours were harvested one week later (see Fig. 3e for dosing regime). Eribulin had no effect on tumour growth in Cre-negative animals compared with vehicle treated animals, but tumour growth was significantly reduced in Eribulin-treated Cre-positive animals (Fig. 3e). Analysis of tumours by immunostaining for blood vessels showed a reduction in intratumoral microvascular density only in sections from Eribulin-treated Cre-positive animals (Fig. 3f). There have been reports that targeting β3-integrin increases the efficiency of drug delivery to tumours (Wong et al., 2015). To test whether this might be driving increased responses in Cre-positive animals, we conducted a similar experiment with doxorubicin, a DNA damaging agent. In these studies, we observed no difference in tumour growth or vessel density when comparing Cre-positive to Cre-negative animals (Fig. 3f). On the whole, the findings suggest β3- integrin depletion in ECs renders them more sensitive to MTAs both *in vitro* and *in vivo.*

The increased sensitivity to destabilising MTAs suggested to us that there is an increased population of stable MTs in β3HET and β3NULL ECs compared with their wild-type counterparts. We explored this premise by exposing ECs to cold temperatures (which destabilises MTs), washing out tubulin monomers (Ochoa et al., 2011), followed by immunolabeling for α-tubulin. We noted elevated stable MTs in both β3HET and β3NULL cells (Fig. 4a). We also measured this biochemically by separately extracting both cold-sensitive (Fig. 4b) and cold-stable (Fig. 4c) MTs from cold-treated cells and Western blotting for α-tubulin. β3HET and β3NULL ECs showed decreased cold-sensitive and increased cold-stable MTs compared with β3WT ECs.

**Figure 4.**
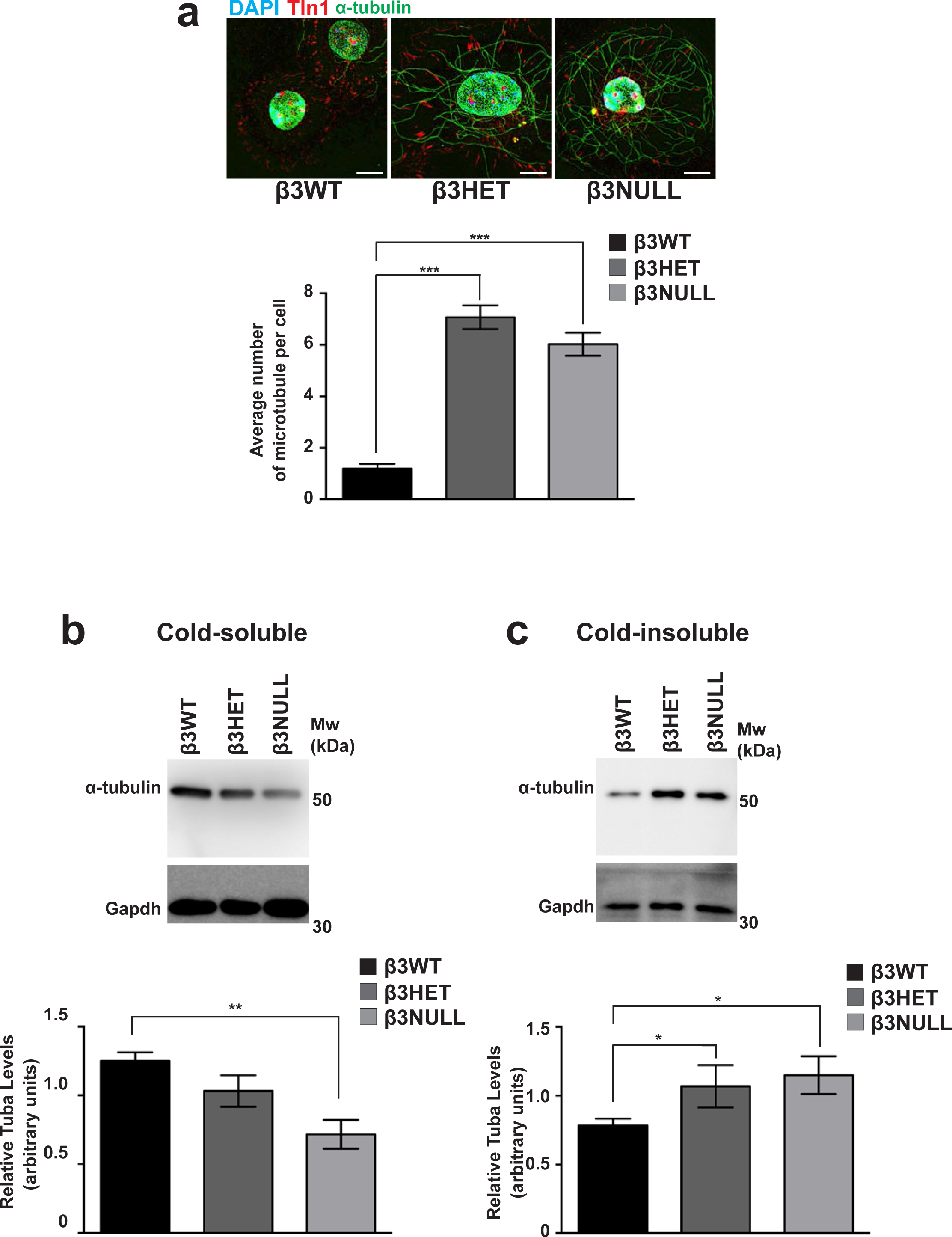
Analysis of microtubule stability in β3WT, β3ΗΕΤ and β3NULL endothelial cells. **a)** *Top* β3WT, β3HET and β3NULL endothelial cells were adhered to fibronectin coated coverslips for 75 minutes at 37°C before being moved to ice for 15 minutes. Soluble tubulin was then washed out using PEM buffer (see materials and methods) before fixing with −20°C methanol. Immunostaining was carried out for α-tubulin (green) and Talin-1 (Tln1-red). DAPI (blue) was used as a nuclear stain. Images shown are representative of the data shown in the bar graph shown below. Scale bar = 20 μm. *Bottom* Bars = mean (±SEM) number of cold-stable microtubules per cell (n=519). **b)** *Top* β3WT, β3HET and β3NULL endothelial cells were adhered to fibronectin for 75 minutes at 37°C before being moved to ice for 15 minutes. Soluble tubulin was then washed out using PEM buffer and soluble material was Western blotted for α-tubulin and Gapdh (as a loading control). Blot is representative of 5 independent experiments. *Bottom* Bars = mean (±SEM) relative cold-soluble α-tubulin levels. **c)** *Top* β3WT, β3HET and β3NULL endothelial cells were adhered to fibronectin for 75 minutes at 37°C before being moved to ice for 15 minutes. Soluble tubulin was then washed out using PEM buffer. The cells were then lysed and insoluble material was Western blotted for α-tubulin and Gapdh (as a loading control). Blot is representative of 4 independent experiments. *Bottom* Bars = mean (±SEM) relative cold-insoluble α- tubulin levels. *= P<0.05; **= P<0.01; ***= P<0.001 in an unpaired, two-tailed t-test.

To gain mechanistic insight into how β3-integrin at FAs might be regulating MT function, we delved deeper into our β3-dependent adhesome data. We noted that Rcc2 clusters with β3-integrin in the β3WT adhesome, but is significantly decreased in that of β3HET ECs. Rcc2 (also known as telophase disk protein of 60 kDa, TD-60) has previously been shown to associate with integrin complexes(Humphries et al., 2009) and to regulate MTs (Mollinari et al., 2003). We therefore examined whether Rcc2 was regulating MT stability in ECs. Knocking down Rcc2 by siRNA in β3WT ECs elicited a significant increase in cold-stable MTs (Fig. 5a). This finding suggested to us that Rcc2 plays a β3-dependent role in regulating MTs in ECs, but does not do so in isolation. We therefore cross-referenced our adhesome data with an Rcc2 pull-down assay performed from HEK-293T cells (Supplementary Table S4) (Williamson et al., 2014). Some obvious potential candidates (e.g. Coronin-1C) were present in both the β3WT and β3HET adhesomes, but at the same level, so were ruled out from further analysis. However, annexin-a2 (Anxa2) co-precipitates with Rcc2 in HEK-293T cells and, like Rcc2, was reduced in the β3HET adhesome. Therefore, we examined whether Anxa2 was also regulating MT stability in ECs via siRNA-mediated knockdown. Like Rcc2 knockdown, even a relatively small (~30%) Anxa2 knockdown in β3WT ECs elicited a significant increase in cold-stable MTs (Fig. 5b).

**Figure 5.**
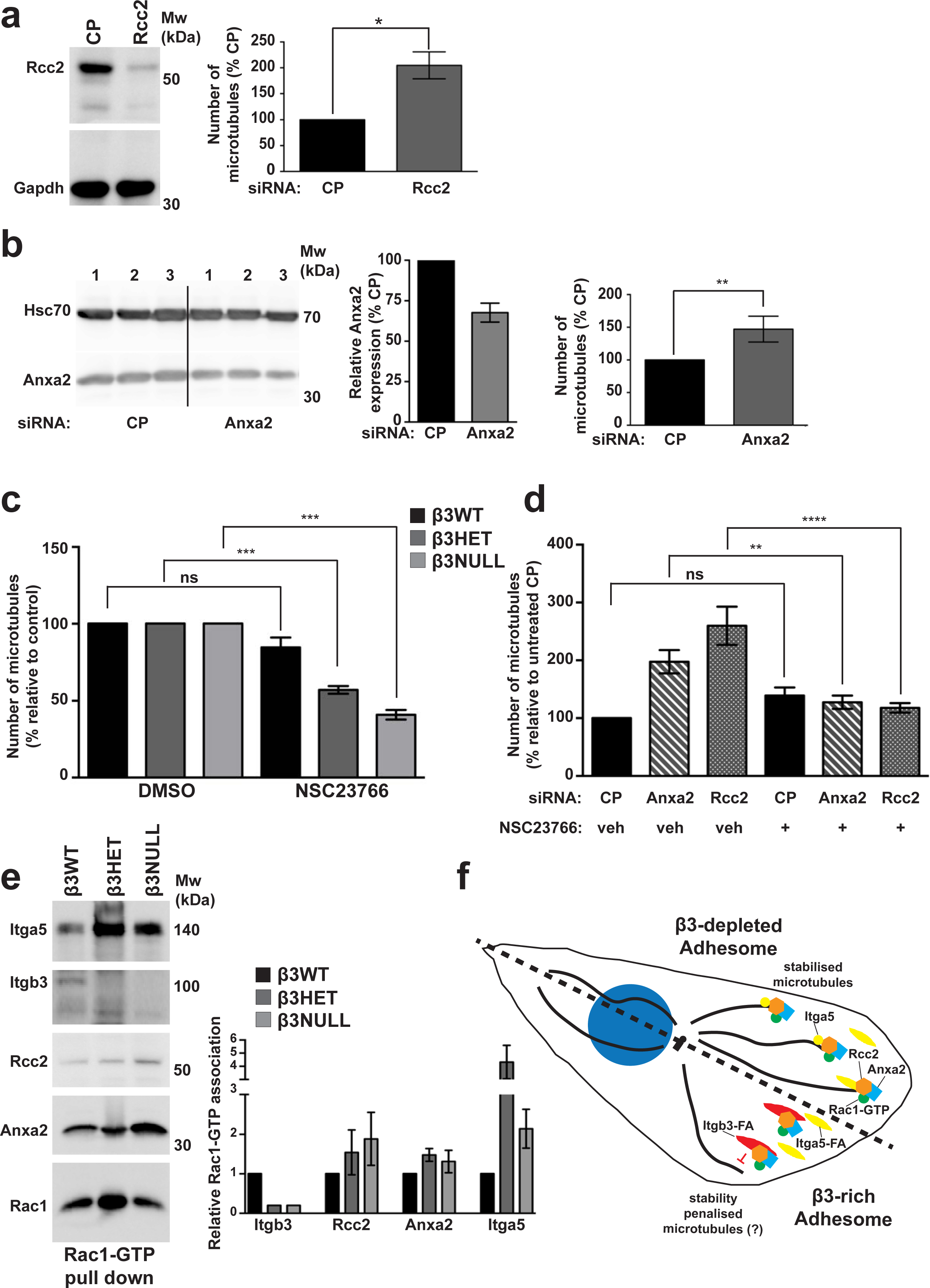
Effects of Rcc2, Anxa2 and Rac1 on microtubule stability in β3WT, β3HET and β3NULL endothelial cells. **a)** β3WT ECs were transfected with control pool (CP) or Rcc2 smart pool siRNA and allowed to recover for 48 hours. They were then adhered to fibronectin coated coverslips for 75 minutes at 37°C before being moved to ice for 15 minutes. Soluble tubulin was then washed out using PEM buffer before fixing with −20°C methanol. Immunostaining was carried out for α-tubulin to allow counting of the number of cold stable microtubules per cell. *Left* Western blot showing representative Rcc2 knockdown. Gapdh is shown as a loading control. *Right* Bars = mean (±SEM) number of cold stable microtubules shown as a percentage relative to CP treated cells (n=224). **b)** β3WT ECs were transfected with control pool (CP) or Anxa2 smart pool siRNA and allowed to recover for 48 hours. They were then adhered to fibronectin coated coverslips for 75 minutes at 37°C before being moved to ice for 15 minutes. Soluble tubulin was then washed out using PEM buffer before fixing with −20°C methanol. Immunostaining was carried out for α-tubulin to allow counting of the number of cold stable microtubules per cell. *Left* Western blot showing representative Anxa2 knockdown in 3 separate samples. *Middle* Bars = mean (±SEM) Anxa2 knockdown shown as a percentage relative to CP treated cells. Samples have been normalised to Gapdh. *Right* Bars = mean (±SEM) number of cold stable microtubules shown as a percentage relative to CP treated cells (n=100). **c)** β3WT, β3HET and β3NULL endothelial cells were adhered to fibronectin coated coverslips for 60 minutes at 37°C before treated with DMSO (control) or 50 μM NSC23766 and incubated at 37°C for a further 15 minutes. Coverslips were moved to ice for 15 minutes. Soluble tubulin was then washed out using PEM buffer before fixing with −20°C methanol. Immunostaining was carried out for alpha-tubulin to allow counting of the number of cold stable microtubules per cell. Bars = mean (±SEM) number of microtubules per cell shown as a percentage relative to DMSO treated controls (n=218). **d)** β3WT endothelial cells were transfected with control pool (CP), Anxa2, or Rcc2 smart pool siRNA and allowed to recover for 48 hours. Cells were then adhered to fibronectin coated coverslips for 60 minutes at 37°C before treated with DMSO (veh) or 50 μM NSC23766 (+) and incubated at 37°C for a further 15 minutes. Coverslips were moved to ice for 15 minutes. Soluble tubulin was then washed out using PEM buffer before fixing with −20°C methanol. Immunostaining was carried out for alpha-tubulin to allow counting of the number of cold stable microtubules per cell. Bars = mean (±SEM) number of microtubules per cell shown as a percentage relative to the CP/veh control (n=100). **e)** *Left* β3WT, β3HET and β3NULL endothelial cells were adhered to fibronectin coated plates for 90 minutes before being lysed in MLB (see Materials and Methods). GTP-Rac1 and bound proteins were extracted from cleared MLB using PAK-1 PBD magnetic beads at 4°C for an hour before being Western blotted for Itga5, Itgb3, Rcc2, Anxa2 and Rac1. Blot is presentative of at least 3 independent experiments. *Right* Bars = mean (±SEM) level of association of the indicated protein with GTP-Rac1, shown relative to β3WT associations. **f)** A schematic model of an endothelial cell adherent to fibronectin that has been “split” down the middle (dotted line) to illustrate conditions of wild-type levels of β3-integrin expression (bottom) or depleted levels of β3-integrin expression (top). When β3- integrin (Itgb3) is expressed at high levels, Rac1-GTP/Rcc2/Anxa2 form a complex that associates with β3-rich FAs. Under these conditions microtubules are either not efficiently stabilised at FAs or they suffer a stability penalty, hence a depletion of microtubules in the β3WT adhesome. However, the loss of β3-integrin allows Rac1-GTP/Rcc2/Anxa2 to localise with α5-integrin. This re-distribution of the complex allows for stabilisation of microtubules, hence their enrichment in the β3-depleted adhesome. Moreover, re-positioning of Rac1 activity means that it plays a role in microtubule-linked EC migration only when β3 is not present in mature FAs. As siRNA mediated knockdown of Rcc2 or Anxa2 in β3WT cells leads to an increase in microtubule stability, we hypothesise that the default association of the complex is with β3-rich adhesions.

Both Rcc2 (Humphries et al., 2009; Williamson et al., 2014) and Anxa2 (Hansen et al., 2002) have been identified as regulators of Rac1, and work by a number of groups has demonstrated that cortical Rac1 activity promotes MT stability (Banerjee et al., 2002; Daub et al., 2001; Wittmann et al., 2004). Because total Rac1 stoichiometry was unchanged when comparing β3WT and β3HET EC adhesomes, we hypothesised that Rcc2/Anxa2-dependent alterations in Rac1 **activity** were responsible for altered MT stability in β3HET and β3NULL ECs. First, we tested the premise that Rac1 plays a differential role in regulating MT stability in β3WT and β3-depleted ECs by testing the effects of the Rac1 inhibitor NSC23766. NSC23766 had no effect on MT stability in β3WT cells, but cold stable MTs in both β3HET and β3NULL ECs were significantly reduced in the presence of the inhibitor (Fig. 5c). We next demonstrated that the increases observed in MT stability upon Rcc2 or Anxa2 knockdown were abrogated in the presence of NSC23766 (Fig. 5d), suggesting that both proteins regulate MT stability in ECs via Rac1.

Rcc2 has been reported to bind directly to the nucleotide-free form of Rac1 and it has been suggested that it functions as a Rac1 guanine nucleotide exchange factor (GEF)(Humphries et al., 2009). Indeed, Rcc2 can guide mesenchymal cell migration by trafficking Rac1 and controlling its exposure to GEFs (Williamson et al., 2014). We therefore tested whether there were differences in Rcc2/Anxa2/active-Rac1 associations between β3WT and β3-depleted ECs. PAK-PBD pull-downs of GTP-bound Rac1 showed co-association of all three proteins in β3WT, β3HET and β3NULL ECs (Fig. 5e), so we concluded that changes in Rac1 activity alone were not responsible for alterations in MT stability in β3-depleted cells. Humphries *et al.* showed that Rcc2 is recruited to α5β1-FN complexes but not α4β1-Vcam1 (vascular cell adhesion molecule-1) complexes in cells expressing both α4- and α5-integrins (Humphries et al., 2009). Thus, we also tested associations between Rcc2, Anxa2 and α5-integrin in β3WT and β3-depleted ECs by PAK-PBD pull-downs. Rcc2, Anxa2, β3- integrin and α5-integrin were pulled down with Rac1-GTP in β3WT ECs. Rcc2, Anxa2 were also pulled down with Rac1-GTP in β3HET and β3NULL ECS, whilst β3-integrin-Rac1-GTP associations were lost and α5-integrin-Rac1-GTP associations were increased (Fig. 5e). Given the stoichiometry of α5-integrin in the β3-depleted adhesome is unchanged compared to the β3WT adhesome (Fig. 2) whilst Rcc2 and Anxa2 levels are decreased, we speculate that a substantial proportion of the observed increase in Rcc2/Anxa2/active-Rac1/Itga5 associations in β3-depleted cells occurs away from FAs, perhaps in recycling endosomes.

In conclusion, by mining the FN-3-integrin EC adhesome, not only have we generated a valuable tool for the integrin and angiogenesis communities, we have also utilised the data to uncover a novel role for β3-integrin in regulating MT function/stability during EC migration. We previously showed that endothelial Rac1 is only required for tumour growth and angiogenesis when β3-integrin is absent (D’Amico et al., 2010), but the underlying mechanism for this observation has remained unclear. Our working hypothesis is that engagement of αvβ3-integrin with FN at mature FAs localises an Rcc2/Anxa2/Rac1 containing complex to these sites, either preventing GTP-Rac1 from participating in MT stability, or actively destabilising MTs (our experiments do not allow us to distinguish between these two possibilities), perhaps by controlling its exposure to GEFs. When αvβ3 is not present, this complex associates with α5β1-integrin instead, where it now has the opposite effect on MTs (Fig. 5f). This re-positioning of Rac1 activity means that it plays a role in MT-linked EC migration only when αvβ3 is not present in mature FAs. There is certainly precedence for β3-integrin regulating spatial distribution of signaling pathway components in cells. For example, we previously showed that β3-integrin plays a role in locally suppressing β1-integin in fibroblasts to promote persistent cell protrusion and migration by regulating interactions between vasodilator-stimulated phosphoprotein (Vasp) and Rap1-GTP-interacting adaptor molecule (Apbb1ip/RIAM) (Worth et al., 2010). Moreover, MTs have recently been shown to target active β1-integrins (Byron et al., 2015). Thus, it will be particularly pertinent to next determine the full composition of the Rcc2/Anxa2/Rac1-GTP complex as many of the proteins that might be suspected to play a role in MT capture (e.g. Clip170 and Clasps) do not appear to be present in the EC adhesome (Fukata et al., 2002); to gain a full picture of how MT stability/FA targeting are regulated in ECs it will be essential to establish how this complex behaves in α5β1-deficient ECs. Finally, whilst our findings reinforce the concept that integrin inhibitors for use as anti-angiogenic agents need rethinking, they nevertheless suggest that once effective αvβ3-integrin antagonists are available (e.g. ProAgio (Turaga et al., 2016)), they may be particularly useful as anti-angiogenic agents when used in combination with already approved MTAs, such as Eribulin.

## Acknowledgements

We thank Maddy Parsons (Kings College London, London, UK) for her gift of the paxillin-GFP construct. A special thanks to both Dr Sophie Akbareian and Peng Liu for their undying enthusiastic and critical support of this project.

## Competing Interests

The authors declare no competing financial interests.

## Author Contributions

SJA designed and performed experiments, analyzed data, and helped write and edit the manuscript. AMG, TSE, and RTJ designed and performed experiments, analyzed data, and helped edit the manuscript. BMK, AA, WJF and BCS performed experiments, analyzed data, and helped edit the manuscript. JGS, KW and MDB provided essential data and helped edit the manuscript. MMM and DRE analyzed data and helped edit the manuscript. SDR designed experiments, performed experiments, analyzed data, and wrote the manuscript.

## Funding

This work was part funded by BBSRC DTP PhD studentships to SJA, RTJ, BMK, and WJF. The work was also part funded by BigC PhD studentships to TSE and AMG and by British Heart Foundation and Breast Cancer Now project grants to SDR and DRE, respectively. The work was also supported by charitable donations from Norfolk Fundraisers, Mrs Margaret Doggett, and The Colin Wright Trust.

## Supplementary Materials and Methods

### Reagents

Unless otherwise stated all chemicals used were obtained from Sigma-Aldrich (Poole, UK). Vascular endothelial growth factor (mouse VEGF-A^164^) was made in house according to Krilleke *et* al.^1^.

### Animals

All animals were on a mixed C57BL6/129 background. Littermate controls were used for all *in vivo* experiments. All animal experiments were performed in accordance with UK Home Office regulations and the European Legal Framework for the Protection of Animals used for Scientific Purposes (European Directive 86/609/EEC).

### Mouse endothelial cell isolation and culture

Mouse lung ECs were isolated from adult mice as per Reynolds and Holdivala-dilke^2^ then subsequently immortalised and cultured as per Ellison *et al.*^*3*^.

### Focal Adhesion Enrichment

Focal-adhesion enrichment was carried out as described in Ellison *et al*.^3^ and Schiller *et al.*^*4*^. A small amount of each focal adhesion sample generated was quality controlled by running a 10% SDS-PAGE gel followed by silver staining (Pierce ^TM^ Silver Stain Kit, ThermoFisher Scientific). Good quality samples were then analysed by western blotting or mass spectrometry.

### Mass Spectrometry (MS)

Mass spectrometry was carried out by the Fingerprints Proteomics Facility, Dundee University, Dundee, UK as per Schiller *et al.*^*4*^. Peptides were identified and quantified using MaxQuant^5^ software using the Andromeda peptide database. To achieve label-free quantitative results, three biological repeats were pooled and each of these pooled samples was analysed via three technical repeats through the spectrometer.

### MS Statistical Analysis

All mass spec analysis was performed using the Perseus^6^ bioinformatics toolbox for MaxQuant. Statistical significance was identified using the Significance Analysis of Microarrays (SAM) method^7^. Unsupervised hierarchical clustering was performed using Perseus’ built in tools. KEGG and GO annotations were obtained from the mouse annotations package via Perseus (downloaded 20/06/2015) and used to identify angiogenesis, cytoskeleton and focal adhesion related genes.

### Random Migration

24 well plates were coated with 10 μg ml^−1^ Fibronectin (FN) in PBS overnight at 4°C and then blocked with 1% BSA for 1 hour at room temperature. 10,000 ECs were seeded per well and allowed to recover overnight. Media was then replaced with media containing one of the following microtubule targeting agents (MTAs): Paclitaxel 5nM, Epothilone B 1nM, Colchine 10μM, Mebendazole 0.4μM, Fosbretablin 0.5μM or Eribulin 1μM (DMSO was used as a control). A phase contrast image was taken of each well every 20 minutes using an inversted Axiovert (Zeiss) microscope for 15 hours at 37°C and 5% CO_2_. The ImageJ plugin MTrackJ^8^ was then used to manually track individual cells and the speed of random migration was calculated.

### Microtubule Stability Assay

Microtubule cold stability assays were carried out as described in Ochoa *et al*.^9^. Briefly:

ECs were seeded per well of a 6 well plate (FN coated/BSA blocked as described earlier) and allowed to adhere for 75 minutes at 37°C before being moved to ice for 15 minutes. Control cells remained at 37°C for the final 15 minutes. Cells were washed with PBS and then 100 μ! of PEM buffer (80 μM PIPES pH 6.8, 1mM EGTA, 1mM MgCl2, 0.5% Triton X-100 and 25% (w/v) glycerol for 3 minutes. A second brief wash was performed with 50μl PEM buffer. All PEM buffer was collected and pooled together with 150μl EB buffer (3% SDS, 60mM Sucrose, 65mM Tris-HCL pH 6.8) at 2X concentration. Remaining material on the plate was then extracted using 300μl of EB buffer. Samples were then used in Western blotting analysis.

Additionally, the same procedure was used on ECs adhered to FN coated/BSA blocked coverslips (acid washed and bake-sterilised before coating). They were treated as above except after PEM washing the slides were immediately immersed in −20°C 100% methanol for 20 minutes. Coverslips were then used in immunocytochemistry analysis.

### *In vivo* tumour growth assay

The syngeneic mouse lung carcinoma cell line (derived from C57BL6 mice) CMT19T was used to grow subcutaneous tumours in β3 fl/fl Tie1Cre positive (and negative littermate control) mice. Under anaesthetic, mice were injected subcutaneously in the flank with 1 × 10^6^ cells. Tumours then grew for 7 days, at which point they were palpable through the skin, before the mice were treated with 0.15mg kg^−1^ Eribulin intravenously once a week for 2 weeks (or 8 mg Doxorubicin kg^−1^ at day 11 and 14 via intraperitoneal injection). After 21 days mice were culled and tumours were excised, photographed and measured for volume using a digital caliper. Tumours were bisected along the midline, fixed overnight in 4% paraformaldehyde, preserved for several days in cryoprotectant (20% sucrose, 2% poly(vinylpyrrolidone) in PBS), embedded in gelatin (8% gelatin, 20% sucrose, 2% poly(vinylpyrrolidone) in PBS) before being snap frozen and stored at −80°C.

### Focal-adhesion and microtubule tracking

1 × 10^6^ ECs were transfected with a GFP-tagged paxillin cDNA expression construct (provided by Dr Maddy Parsons, King’s College London, London, UK) by nucleofection. Cells were allowed to recover overnight before a fraction were seeded on FN coated/BSA blocked coverslips (acid washed and baked before coating) and adhered for 3 hours. Cells were then treated with 100nM SiRTubulin (Cytoskeleton Inc CY-SC002) and 1μM Verapamil overnight. Coverslips were imaged individually on an Axiovert (Zeiss) inverted microscope where one image of a GFP positive cell was taken every minute for 30 minutes at 37°C and 5% CO_2_ in green and far-red channels. During imaging media was replaced with Phenol-red free OptiMEM® + 2% FBS containing 100nM SiRTubulin and 1μM Verapamil. The total area of adhesive fronts was assessed by measuring the growth of paxillin-GFP positive areas between the 1^st^ and 30^th^ image and then the number of microtubules that entered the adhesive front over 30 minutes were counted.

### Western Blotting

For western blot analysis of total tubulin levels. ECs were seeded at 750,000 per well of a FN coated/BSA blocked 6 well plate and allowed to adhere for 90 minutes before being lysed in EB buffer. For the microtubule stability assay and focal adhesion enrichment samples were prepared as above. 20μg from each sample was loaded onto 10% polyacrylamide gels then transferred to a nitrocellulose membrane and incubated for 1 hour in 5% milk powder in PBS with 0.1% Tween 20 (PSBTw) followed by overnight incubation in primary antibody diluted

1:1000 in 5% BSA in PBSTw at 4°C. Primaries used were against integrin beta 3 (Cell Signalling 4702), alpha-tubulin (Abcam 7291), Gapdh (Abcam 9484), Rcc2 (Abcam 70788), Hspa1a (clone B-6 Santa Cruz Biotechnology), Anxa2 (Abcam 41803), and Itga5 (Cell Signalling 4705). The membranes were then incubated with the appropriate horseradish peroxidase (HRP)-conjugated secondary antibody (Dako) diluted 1:2000 in 5% milk in PBSTw for 1 hour at room temperature. The blot was visualised using Piece^®^ ECL Western Blotting Substrate kit (ThermoFisher) and chemiluminescence detected on a Fujifilm LAS-3000 darkroom (Fujifilm UK Ltf, Beford, UK).

### Immmunocytochemistry

ECs were seeded onto FN coated/BSA blocked coverslips and adhered for 90 minutes before being washed with PBS and immersed in −20°C methanol for 20 minutes. Alternatively, cells were prepared as per the microtubule stability assay protocol above. Coverslips were then washed with PBS, blocked for 10 minutes at room temperature with 0.5% BSA, 1% goat serum in PBS with 0.25% Triton X-100 and incubated with primary antibody diluted 1 in 250 in PBS for 1 hour at room temperature. After subsequent PBS washed the coverslips with incubated with the appropriate goat raised Alexa-Fluor® conjugated secondary antibody (Invitrogen) diluted 1 in 500 in PBS. Coverslips were washed again in PBS before being mounted into slides using Prolong^®^ Gold with DAPI (Invitrogen). Primaries used alpha-tubulin (Abcam 52866), paxillin (Abcam 32084) and talin (Sigma T3287). Nrp1 (R&D Systems AF566) staining was carried out as above except with donkey serum, and donkey raised Alexa-Fluor® secondary.

Simultaneous Phalloidin (ThermoFisher A12380) and alpha-tubulin staining was carried out using PHEMO fixation ^10^.

### Immunohistochemistry

5 μm cryosections were prepared from frozen tumours and stained as described previously^3^. Images were acquired on an Axioplan (Zeiss) epifluorescent microscope. Additionally, scans of complete sections were achieved using the AxioVision MosaiX plugin. Tissue area and vessel counts were obtained using ImageJ, as described in Ellison *et al.*^*3*^.

### siRNA knockdown

Knockdowns of Rcc2 and Anxa2 were achieved using 3μg of Dharmacon ON-TARGETplus SMARTpool siRNA (control pool used as knockdown control) per 1 × 10^6^ ECs in an Amaxa Nucleofector II (T-005 setting). Cells were allowed to recover for 48 hours to allow knockdown to take effect.

### Active Rac1 Pulldown

6 × 10^6^ ECs were seeded onto FN coated/BSA blocked 10 cm plates and allowed to adhere for 90 minutes. Rac1 Activation Magnetic Beads Pulldown Assay kit (Millipore 17-10393) was then used per manufacturer’s instructions. Pull-down material was then loaded directly onto a gel for western blotting.

